# DeepVul: A Multi-Task Transformer Model for Joint Prediction of Gene Essentiality and Drug Response

**DOI:** 10.1101/2024.10.17.618944

**Authors:** Ala Jararweh, My Nguyen Bach, David Arredondo, Oladimeji Macaulay, Mikaela Dicome, Luis Tafoya, Yue Hu, Kushal Virupakshappa, Genevieve Boland, Keith Flaherty, Avinash Sahu

**Affiliations:** Comprehensive Cancer Center, The University of New Mexico; Department of Computer Science, The University of New Mexico; Department of Biomedical Engineering, The University of New Mexico; Department of Medicine, Massachusetts General Hospital Cancer Center, Harvard Medical School; Department of Mathematics and Statistics, The University of New Mexico

## Abstract

Despite their potential, current precision oncology approaches benefit only a small fraction of patients due to their limited focus on actionable genomic alterations. To expand its applicability, we propose DeepVul, a multi-task transformer-based model designed to predict gene essentiality and drug response from cancer transcriptome data. DeepVul aligns gene expressions, gene perturbations, and drug perturbations into a latent space, enabling simultaneous and accurate prediction of cancer cell vulnerabilities to numerous genes and drugs. Benchmarking against existing precision oncology approaches revealed that DeepVul not only matches but also complements oncogene-defined precision methods. Through interpretability analyses, DeepVul identifies underlying mechanisms of treatment response and resistance, as demonstrated with BRAF vulnerability prediction. By leveraging whole-genome transcriptome data, DeepVul enhances the clinical actionability of precision oncology, aiding in the identification of optimal treatments across a broader range of cancer patients. DeepVul is publicly available at https://github.com/alaaj27/DeepVul.git.

## 1. Introduction

Precision oncology seeks to match patients with the most effective drugs. Current approaches benefit only 10% of cancer patients due to their narrow focus on a small number of actionable genomic alterations [14, 2, 20]. Consequently, over 90% of cancer patients lacking actionable alterations in their tumors are ineligible for these therapies. To expand the coverage of precision oncology beyond this 10%, “avatar models” have been proposed. These models involve creating cell lines, xenografts, organoids, or ex vivo systems from tumor biospecimens from a patient, which can then be tested with numerous drugs or gene knockouts to find the most optimal treatment [8, 13]. The time required to develop these experimental models impedes their clinical adoption and practical application. To address this, researchers are now mapping targetable cancer vulnerabilities in a large number of existing cell lines and mouse xenografts, aiming to translate insights gained from these experimental approaches, including drug treatments and gene essentiality screens, into patient care.

Gene essentiality in cancer cells refers to the degree to which the survival and proliferation of the cancer cell depends on the function of a specific gene [32, 11, 23]. It is quantified by the change in growth rate resulting from the knockout or knockdown of that gene, indicating how critical the gene is to the cell’s viability. Mapping essential genes reveals specific cancer cell vulnerabilities, providing potential therapeutic targets. By identifying selective gene knockouts that can kill specific tumor cells without affecting normal cells, researchers can design drugs targeting these genes, thereby increasing the coverage of precision oncology beyond the few currently known actionable genes. Experimental approaches such as CRISPR knockout screens reveal essential genes on a genome-wide scale [11, 31]. For instance, the DepMap project has mapped essential genes in thousands of cancer cell lines. [28, 6].

Complementing experimental approaches, computational approaches have been proposed for predicting gene essentiality. This includes traditional machine learning approaches like Support Vector Machines (SVM) using features from nucleotide sequence data and protein-protein interaction (PPI) networks, which require manual feature selection and domain knowledge [9, 36, 33]. Deep learning approaches have also been adopted for essentiality prediction because of their ability to perform feature extraction and engineering without human intervention [35, 36, 12].

However, gene essentiality varies dramatically across cancer cell lines, making it highly context-specific [28]. This context-specificity raises questions about how findings in cell lines can be translated to patient tumors. The relationship between the cancer genetic landscape and its vulnerabilities is non-linear and complex, necessitating advanced modeling for accurate predictions. Current methods fail to capture this context-specificity accurately, preventing us from effectively assessing the impacts of gene knockouts on cancer cells without modeling cancer as a system.

Here, we posit that mapping gene expression to cancer vulnerability could enable the identification of optimal treatments for cancer patients from their tumor transcriptome. We propose DeepVul, a multi-task transformer-based model that aligns gene expression and essentiality into a latent space, allowing the joint prediction of drug response and gene essentiality. Our framework jointly predicts the vulnerability of cancer cells to thousands of genes and their responses to thousands of drugs simultaneously. Our interpretability framework also identifies underlying response mechanisms, as demonstrated for the BRAF oncogene. By leveraging whole-genome transcriptome data, our approach aims to expand the clinical actionability of precision oncology, identifying optimal treatments for cancer patients.

## 2. Related Work

There are two types of gene essentiality: universal and context-specific. Universal essential genes are vital for the survival of an organism across all contexts, external conditions, or environments, and are typically involved in critical cellular processes (Appendix C). Context-specific essential genes refer to those genes that are vital for a cell’s survival in specific environmental conditions, and targeting these genes would kill specific cells without affecting others. Therefore, context-specific essential genes are of particular interest in personalized medicine, as they may be targets for drug development, especially for diseases like cancer. In this study, we focus on context-specific essential genes.

### Context-Specific Gene Essentiality

DeepHE, [36], a neural network, classifies essential genes in human cancer cell lines using sequence and PPI data; however, assumes that genes are universally essential across cancer cell lines. FluxGAT, [25] is a GNN-based approach for predicting gene essentiality directly from graphical representations of flux sampling data. DeepDEP, [3], a deep learning model, predicts essential genes for cancer cell proliferation using integrative genomic profiles.

### Multi-Task Learning

Multi-task learning refers to the joint training of a single machine learning model on multiple tasks. To improve retinal blood vessel segmentation, a novel encoder-decoder architecture leveraging multi-task learning has been proposed, [16]. Clustering-Constrained-Attention Multiple-Instance Learning (CLAM) is another multitask framework for whole slide image (WSI) processing, which utilizes attention-based learning for predicting multiple tasks, [18]. Another multitask architecture, DeX [26], applies a transformer-based approach to model interactions between input features using the attention mechanism. Given the interactive nature of genes, this architecture is effective for genomics applications. Our proposed method, DeepVul, draws inspiration from this architecture.

## 3. Methodology

We propose DeepVul, a multi-task transformer-based model that connects gene expression and perturbations into a latent distribution, inspired by DeX [26]. The paper mainly discusses the significant challenge of predicting throughput in real-time communication in extreme conditions to enable proactive network management. The proposed model comprises three major components, feature selection, feature extraction via a transformer-based framework, and multi-task predictions to predict regular throughput, traffic peaks, traffic valleys, etc. We draw an analogy of predicting multiple gene essentiality for an expression profile and predicting different throughput tasks from packet-level information from network traffic. However, we expand upon their model by integrating a few modifications tailored to the essentiality task. Our framework comprises three main components(Figure 1): feature transformation, feature extraction, and fine-tuning. During feature transformation and feature extraction, the model learns a latent space of gene expression relative to gene essentiality, and perturbation data, by utilizing a multi-task loss function that reconstructs essentiality data from expression. Next, we fine-tune the latent space by training a second multi-task objective that predicts drug responses by minimizing each drug’s Mean Squared Error (MSE). Finally, we evaluate the model’s performance across different predictive tasks on unseen cohorts of essentiality data.

**Figure 1:**
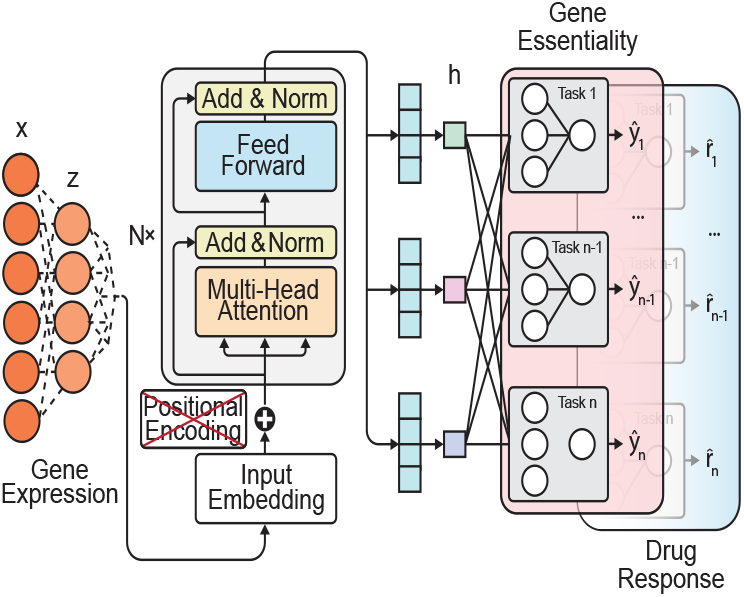
DeepVul architecture: a multi-task deep learning model for predicting gene essentiality and drug response from gene expression using latent space and sequential fine-tuning.

### 3.1. Data Preparation

We utilize the open-source gene essentiality, gene expression, and drug response datasets from the DepMap Achilles project [5] (Appendix C). Because the experiment measured drug response data in the same cell lines, it presents rich information for finding the latent space described below.

We build a multimodal model that integrates the three modalities to identify drug treatments by matching them to the gene expression profiles of specific tumors. From the dataset, we retained cell lines with both gene expression data, and either essentiality or drug response data available. Our study focuses primarily on highly variable and actionable genes and drugs that exhibit significant variability across cells. We utilize the Variance-stabilizing transformation (VST) method to identify the top 1000 highly variable genes and drugs from each dataset [21, 10].

### 3.2. DeepVul architecture

Our goal is to map gene expression profiles of cancer cells to their essentiality and drug vulnerabilities. For this, we train an encoder to encode gene expression into a latent space and use the space to predict essentiality or drug response using decoders. The encoder is shared across all tasks, while decoders are task-specific. Below we describe the architecture and training of encoders and decoders.

#### Encoder: Finding a Latent Space

Input: Gene expression; Output: Latent space that connects expression and essentiality. Initially, we reduce the dimensionality of the gene expression data, **x** ∈ ℝ^*n*^ through a series of linear transformations, resulting in **z** ∈ ℝ^*d*^. We then utilize a stack of transformer layers to process the transformed input, aiming to capture complex relationships among genes:

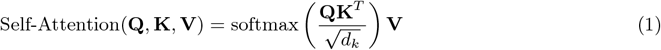

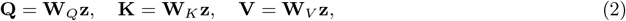

where (**W**_*Q*_, **W**_*K*_, **W**_*V*_) are learnable weights, and *d*_*k*_ is a dimension of the key vector used for scaling. The output, *h*, is then pooled and passed through a feedforward network:

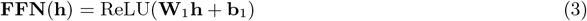

Since positional information is irrelevant in gene expression data, we eliminate positional encoding from the transformer architecture. Attention captures dependencies between genes based on their expression profiles without considering proximity.

#### Essentiality Decoder: Learning Multi-gene Essentiality Predictors

The decoder comprises independent pipelines (MLP layers) for each gene, allowing tunable layers specific to each gene while maintaining the same latent space (Fig 1). Each gene’s pipeline operates independently, unaffected by the pipelines of other genes. The encoder and essentiality decoder are learned jointly leveraging a multi-task objective that fine-tunes the latent space and predicts the essentiality of every gene at a time as:

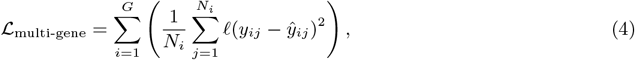

where *G* is the number of genes in the essentiality dataset, *N*_*i*_ is the number of samples for the i-th gene, and *ŷ*_*ij*_ = MLP_*i*_(*Z*) is the prediction for the i-th gene by feeding the latent space to a multi-layer perceptron.

#### Drug Decoder: Learning Multi-drug Response Predictor through Fine-tuning

The drug decoder also comprises an independent pipeline for each drug, similar to the essentiality decoder. The latent space learned by predicting gene essentiality from expression profiles is then fine-tuned to predict drug responses using multi-task objective layers as:

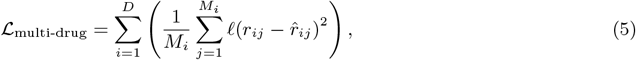

where *D* is the number of drugs in the drug response dataset, *M*_*i*_ is the number of samples for the *i*-th drug, and 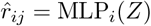 is the prediction for the *i*-th drug by feeding the latent space to a multi-layer perceptron. The drug response pipeline leverages the weights of the latent space to initially create the latent space for the multi-task drug layers. Subsequently, the weights of the latent space can be either fine-tuned or frozen while the multi-task layers are always trainable.

### 3.3. SHAP Analysis

**SH**apley **A**dditive Ex**P**lanations (SHAP) is a method of assigning contributions of inputs to a black box model’s output based on game theory [19]. For a given input, the contribution of each feature sums to the difference between the model output and the model expected value. We calculate the SHAP values for all 19,193 gene expression inputs as they contribute to the model output of gene essentiality. We then use a correlation analysis [15] in which the Pearson correlation between the SHAP contribution and the expression value for a given gene is used to rank genes with the highest and lowest correlations.

## 4. Results

### 4.1. Predictions of Essential Genes in Cancer Cell Lines

We benchmarked DeepVul against multiple baselines (refer to Appendix E and Appendix D for baselines and training details). Evaluating the flattened test data by ignoring cell lines and genes demonstrates that DeepVul shows a high correlation between predicted and actual essentiality across genes and cell lines (Figure 2a). The baseline Autoencoder-NN model also performs similarly in terms of correlation (Figure 2b). However, when we calculated the correlation for each gene separately, DeepVul outperforms the baselines including Autoencoder-NN (Figure 2c). This suggests that when the essentiality prediction task was assessed one gene at a time, predicting how the essentiality of a given gene varies across cell lines, the task becomes challenging. All models including DeepVul showed zero or often negative correlations for many genes. Nonetheless, DeepVul predicted essentiality for 46% of genes with a correlation greater than 0.3.

**Figure 2:**
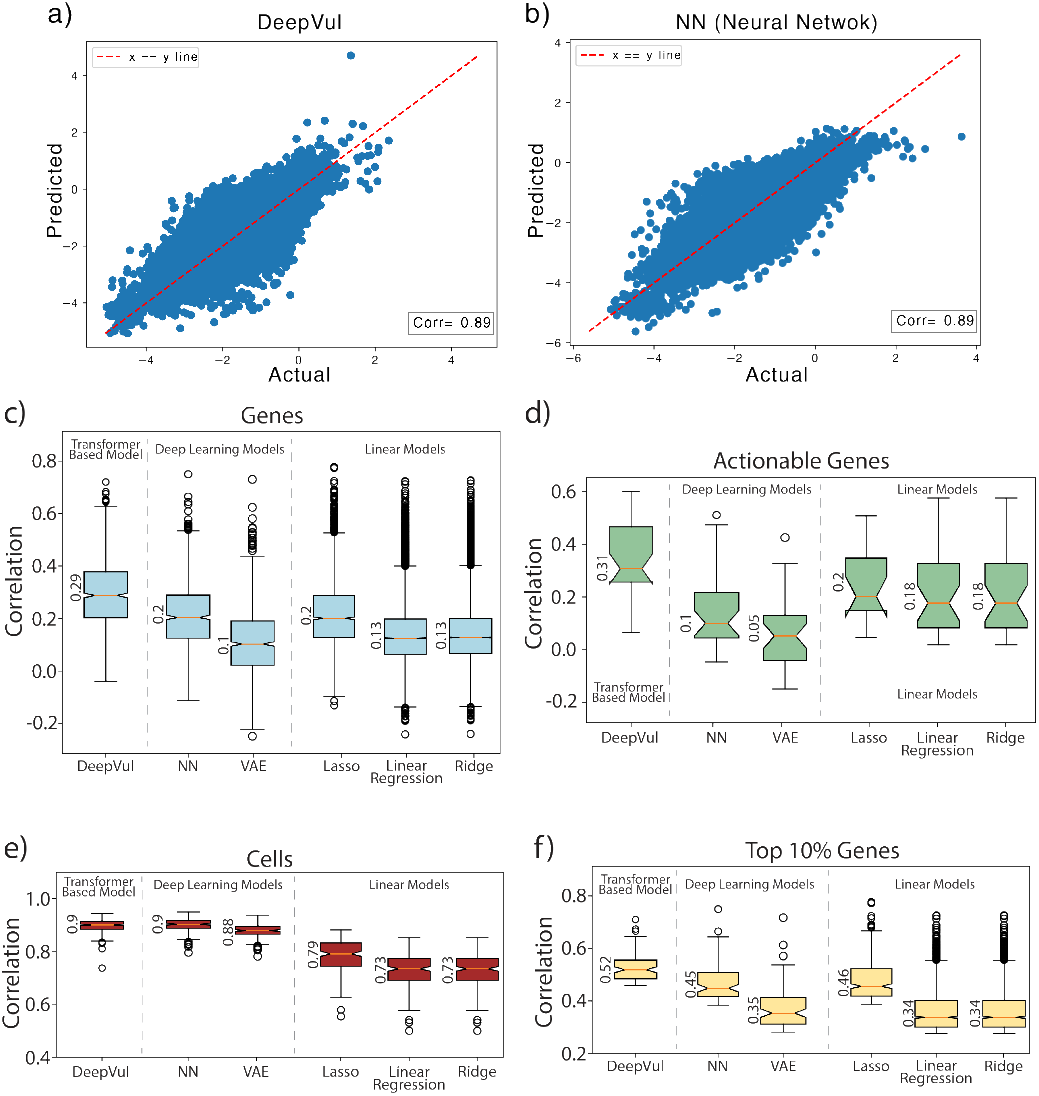
Evaluation of DeepVul compared to various baseline methods in predicting gene essentiality. a,b) Scatter plot shows the predicted vs. actual values of DeepVul and the neural network baseline. c,d,e) Bar charts illustrate a detailed evaluation of our model to the baselines on various prediction categories such as genes(c), actionable genes (d), and cells(e).

The performance gap between DeepVul and the baselines increased when we analyzed important genes known to be actionable (Figure 2d). Calcu-1 lating the correlation between predicted and actual essentiality for each cell line separately, DeepVul and Autoencoder-NN showed similar performance (Figure 2e). DeepVul matched or outperformed the baseline models, particularly in predicting the essentiality of a single gene and how it varies across cell lines.

#### Generalizability to New Cohorts

Model generalizability to new cohorts is a primary challenge widespread across biomedical applications of AI. We evaluated DeepVul’s ability to generalize predictions to an independent cohort without additional training. Using data from the Sanger Institute, we assessed DeepVul’s performance, which was initially trained on DepMap essentiality data from the Broad Institute.

DeepVul’s performance on the Sanger Institute data showed a similar correlation (Figure 3a,b), indicating that DeepVul’s predictions could generalize to new datasets without requiring further training. In contrast, the baseline model failed to generalize effectively (Figure A.8).

**Figure 3:**
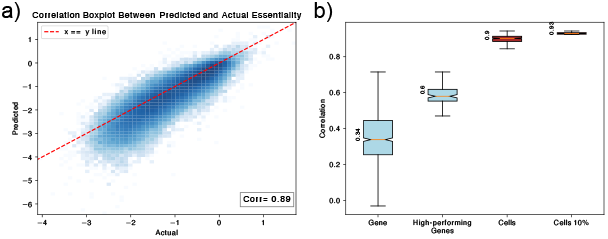
Evaluation of DeepVul predictions on Sanger cohort. a) Scatter plot shows the predicted vs actual gene essentiality values on the Sanger dataset. b) The box-plot illustrates the predictive performance across gene and cell lines.

### 4.2. Predicting Drug Response via Transfer learning

We next evaluate DeepVul’s transfer learning capability by assessing its ability to predict responses to multiple drugs. We test two transfer learning paradigms: one with a frozen expression encoder (referred to as freeze-shared) and another with a tunable expression encoder fine-tuned for drug response (referred to as tuned-shared). Surprisingly, the model with the frozen encoder outperforms the tunable encoder in predicting drug responses (Figure 4).

**Figure 4:**
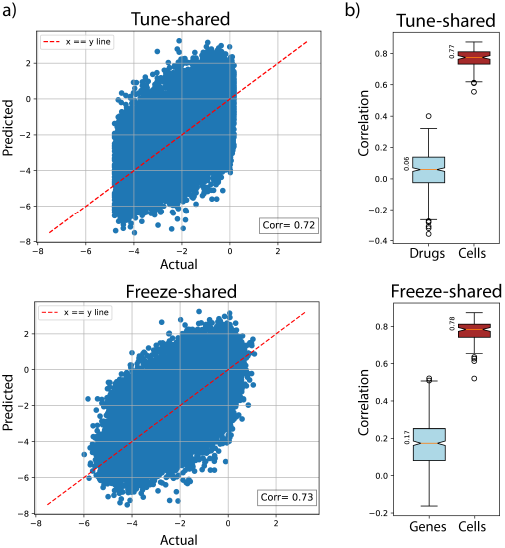
Comparison of drug response prediction methods: Fine-tuning or Freezing the latent shared space. (a) The correlation between the actual and predicted drug response values when the latent shared space is either fine-tuned (left) or frozen (right). (b) The Drug and cell-wise correlation of the two training paradigms.

This outcome may be likely due to drugs targeting a smaller set of pathways compared to gene essentiality, which is measured genome-wide. As a result, gene essentiality might capture cancer at a systems level more comprehensively than drug response. Additionally, the limited training data available for drug response may explain why fine-tuning the parameters does not enhance performance for drug prediction. This finding aligns with transfer learning paradigms in the imaging field, where pre-trained models like ResNet and VGG show no performance gain when fine-tuned on small datasets [4]. Fine-tuning is recommended only when large training datasets are available.

### 4.3. Benchmarking Against Current Precision Approaches

We benchmarked DeepVul against current actionable-mutation-based precision oncology approaches, which recommend drugs based on the presence of single actionable mutations in tumors. For instance, EGFR tyrosine kinase inhibitors (EGFR-TKIs) are recommended as the first-line treatment for patients with EGFR-mutant non-small cell lung cancer [7]. Similarly, BRAF inhibitors are the first-line treatment for patients with BRAF-mutant melanoma, particularly those with the BRAF V600E mutation [27].

DeepVul’s predictions were compared to these established precision oncology approaches. Specifically, DeepVul predicted higher essentiality of the BRAF gene in BRAF-mutant cell lines than in those without BRAF mutations (Figure 5a). The DepMap project defines an essentiality threshold of −1 to indicate sensitivity [28]. We used this threshold for predicted essentiality to identify cell lines sensitive to BRAF targeting and compared the experimental essentiality of our predicted sensitive cell lines with BRAF-mutant cell lines. The experimental essentiality of our predicted sensitive cell lines was slightly lower than that of the BRAF-mutant cell lines (Figure 5b), although the difference was not significant due to the small number of cell lines with predicted essentiality of less than -1 in our test set. This suggests that DeepVul can identify cell lines sensitive to BRAF targeting similar to existing precision oncology.

**Figure 5:**
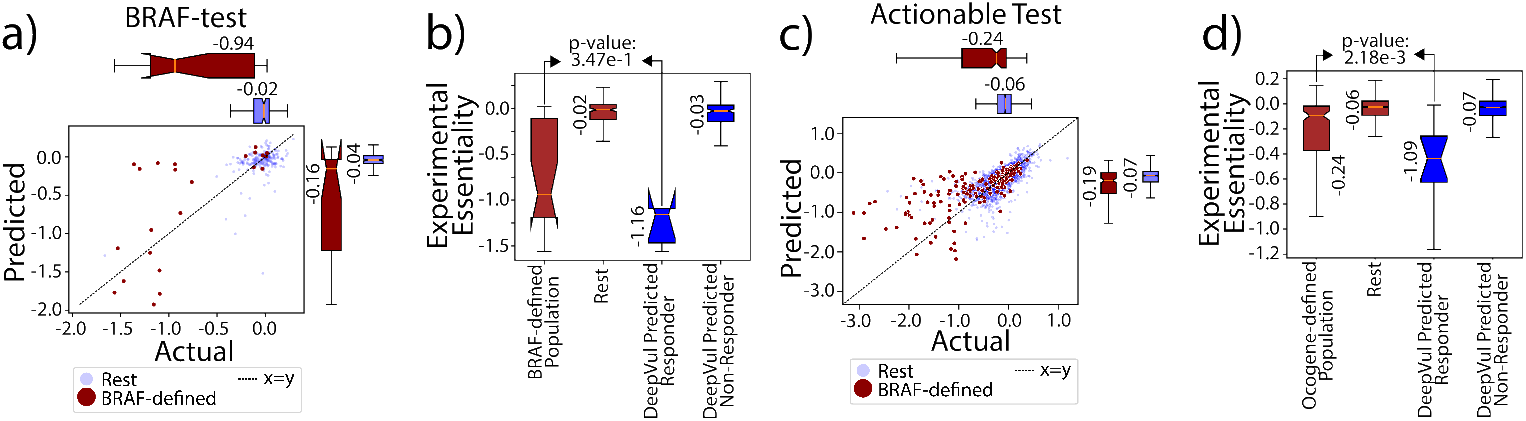
Comparative analysis between DeepVul and other clinical decisions regarding actionable genes. a) Predicted vs. actual essentiality values for BRAF-defined (red) and other cell lines (blue). Boxplots on the side summarize the essentiality difference between BRAF-defined vs other. b) Experimental essentiality of cell lines predicted as responders and non-responders by DeepVul and the corresponding oncogene-based clinical decisions. Wilcoxon P-value for the difference. c) Similar to (a), but for all actionable genes instead of just BRAF. d) Similar to (b), but for all actionable genes instead of just BRAF.

We extended this analysis to other 32 actionable mutations, such as PIK3CA and EGFR [30]. Similar to the BRAF analysis, DeepVul predicted higher essentiality for mutant cell lines, indicating its ability to capture current oncogenic-based populations (Figure 5c). For each actionable mutation, we predicted sensitive cell lines based on the threshold of predicted essentiality of −1. DeepVul-predicted sensitive cell lines showed significantly lower experimental essentiality than those selected based on oncogenic-defined populations (Figure 5d). This suggests that DeepVul can identify target populations for current targeted therapies, complementing existing precision oncology approaches.

### 4.4. SHAP and LISA interpretability identify resistance mechanism

SHAP values provide the relative contribution of a given input feature. Gene expressions are often highly correlated due to co-regulation by transcription factors. To decouple these co-expression patterns, we identified genes whose SHAP values were correlated with their expression levels and identified their transcription factor regulators using LISA [22]. We applied this procedure to the essentiality of BRAF (Figure A.7).

For instance, in the case of positive correlation, a cell with higher expression levels of APOL1 is likely to depend on BRAF1 as an essential gene. The transcription factor regulators of these genes are listed in Table B.1. Among them, STAT3 is the third most likely transcription factor regulator. Inhibition of STAT3 is known to enhance sensitivity to BRAF inhibitors [37], while activation of this pathway is associated with BRAF inhibitor resistance [17]. This analysis suggests that the interpretability framework of DeepVul could identify treatment response and resistance mechanisms.

## 5. Conclusion

We introduced DeepVul, a multi-task transformer-based model designed to predict gene essentiality and drug response from gene expression profiles. Our approach aligns gene expressions, gene perturbations, and drug perturbations into a latent space, enabling simultaneous predictions of cancer cell vulnerabilities to numerous genes and drugs. For gene essentiality prediction, DeepVul matched or outperformed the baseline models and generalized to new datasets without requiring further training. Using BRAF as a use case, we demonstrated that DeepVul can identify cell lines sensitive to targeting hundreds of genes with similar accuracy to existing precision oncology approaches currently available for only a few actionable genes. Furthermore, its interpretability framework could identify treatment response and resistance mechanisms. Thus, DeepVul can aid in identifying target populations for current targeted therapies and complement existing precision oncology approaches.

## Appendix A. Figures

**Figure A.6:**
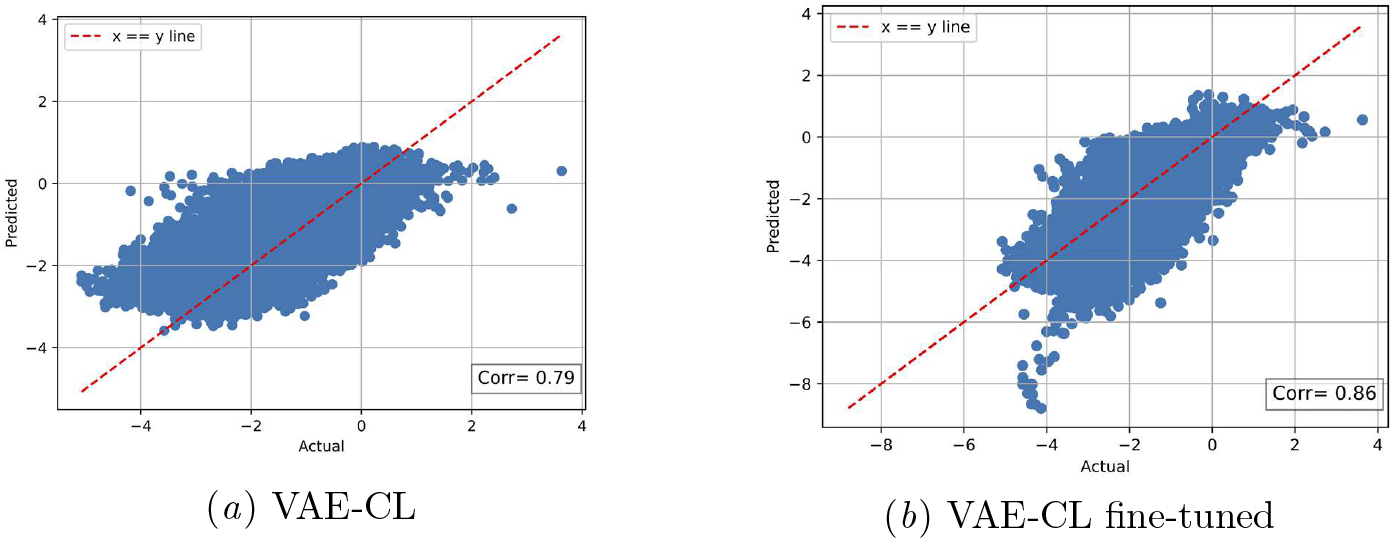
The prediction and actual essentiality values of the baseline VAE-CL before and after fine-tuning.

**Figure A.7:**
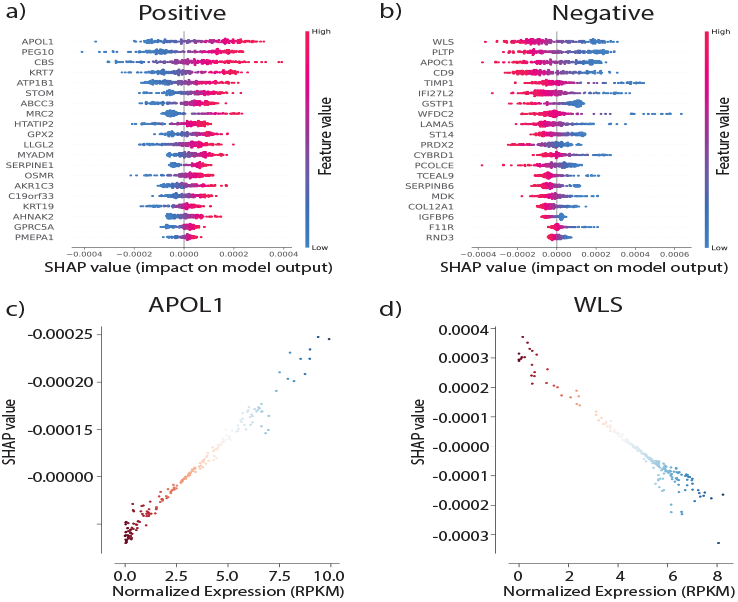
SHAP-expression correlation analysis for BRAF1. **a)** Shap summary plot showing SHAP value (x-coordinate) and Normalized Expression value (color) for the top 20 genes (y-axis) with the highest positive Pearson correlation between SHAP and Expression values, where each point represents one sample in the test set. **b)** Summary plot for highest negative correlations. **c)** Scatter plot of SHAP value (y-axis) vs. Normalized Expression value (x-axis) for the gene with the highest positive correlation (APOL1) shown in *a*. **d)** Scatter plot for the highest negative correlation (WLS) shown in *b*.

**Figure A.8:**
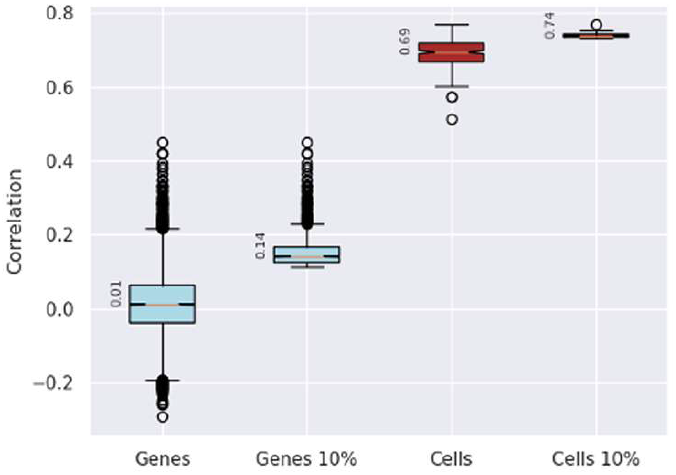
The performance of the VAE-CL fine-tuned in predicting Sanger cohort. The figure reports the Spearman correlation coefficient of the predicted and actual essentiality values across genes and cell lines.

## Appendix B. Tables

**Table B.1:** LISA-Inferred Transcription Factor Regulators of Important Genes Identified Using SHAP Interpretability.

| TF | Rank | Sample p-value |  |  |  |  |
| --- | --- | --- | --- | --- | --- | --- |
|  |  | 1st | 2nd | 3rd | 4th | 5th |
| HMG20A | 1 | $7.09 \times 10^{-13}$ | $2.44 \times 10^{-1}$ | N/A | N/A | N/A |
| FOS | 2 | $2.31 \times 10^{-10}$ | $5.97 \times 10^{-10}$ | $7.15 \times 10^{-10}$ | $4.68 \times 10^{-9}$ | $7.23 \times 10^{-8}$ |
| STAT3 | 3 | $1.13 \times 10^{-9}$ | $1.24 \times 10^{-8}$ | $2.23 \times 10^{-6}$ | $1.03 \times 10^{-5}$ | $1.26 \times 10^{-5}$ |
| JUN | 4 | $1.23 \times 10^{-9}$ | $3.76 \times 10^{-9}$ | $8.14 \times 10^{-8}$ | $1.21 \times 10^{-7}$ | $1.02 \times 10^{-5}$ |
| YAP1 | 5 | $2.70 \times 10^{-9}$ | $4.45 \times 10^{-8}$ | $1.76 \times 10^{-5}$ | $2.48 \times 10^{-3}$ | $4.17 \times 10^{-3}$ |
| TEAD1 | 6 | $4.47 \times 10^{-9}$ | $6.95 \times 10^{-8}$ | $4.93 \times 10^{-5}$ | $2.45 \times 10^{-4}$ | $1.26 \times 10^{-2}$ |
| TP53 | 7 | $6.42 \times 10^{-9}$ | $4.39 \times 10^{-6}$ | $6.39 \times 10^{-5}$ | $1.82 \times 10^{-4}$ | $6.17 \times 10^{-4}$ |
| RELA | 8 | $2.30 \times 10^{-8}$ | $6.36 \times 10^{-8}$ | $2.86 \times 10^{-4}$ | $3.05 \times 10^{-4}$ | $3.31 \times 10^{-4}$ |
| TEAD4 | 9 | $3.58 \times 10^{-8}$ | $1.22 \times 10^{-5}$ | $1.86 \times 10^{-5}$ | $2.35 \times 10^{-5}$ | $4.12 \times 10^{-5}$ |
| SMAD4 | 10 | $4.74 \times 10^{-8}$ | $9.00 \times 10^{-3}$ | $1.53 \times 10^{-1}$ | $2.34 \times 10^{-1}$ | $5.69 \times 10^{-1}$ |

## Appendix C. Related Work

### Universal Gene Essentiality

DeeplyEssential, a multi-layer perceptron (MLP) based network, predicts universal essential genes in microbes using features from DNA and protein sequences [12]. EPGAT, a Graph Neural Network (GNN) based approach predicts universal essential genes from protein-protein interaction networks and multiomics data [24]. EPGAT was benchmarked for four organisms, including humans.

### Experimental Approach

Gene essentiality can be measured using CRISPR-Cas9 in pooled or arrayed screens. Pooled screens deploy many gRNAs simultaneously, enabling high-throughput, cost-effective screening of thousands of genes. Arrayed screens use fewer gRNAs per gene, offering higher resolution and specificity [1]. The DepMap Achilles project aims to find essential genes in cancer cell lines via genome-scale, pooled CRISPR knockout screening, by knocking out genes in various cancer cells and measuring the effect on cell proliferation and survival [5]. DepMap also measures response to different therapeutic compounds in various cancer cell lines [28].

## Appendix D. Training Details

Both models were trained using a comprehensive set of hyperparameters to optimize the model performance. The model utilizes the Adam optimizer of the learning rate set to 5*e*^*−*4^, a batch size of 40, ReLU activation, 20 epochs for each model, and a dropout rate of 0.2. For the transformer layers, we use 2 transformer encoder layers with 4 heads, a hidden state of 1000 for the linear transformation layer, and a 2048 dimension for the feedforward layer. All experiments were performed on an NVIDIA A100 PCIe GPU with 80GB of memory with an estimated time of 1.2 seconds per epoch. We have not performed any hyper-parameter search experiments to find the parameters and the parameters chosen are based on the recommendations found in the original transformer paper as well as computational upper bounds [29]. Yet, we expect the model performance to be further increased if a parameter search is performed.

## Appendix E. Baseline Models

To evaluate the performance of our proposed DeepVul framework, we compare it against several baseline models of different complexity. First, we attempt to approach the problem using linear regressors found in the Sklearn library such as Linear Regression ^1^, Lasso regressor ^2^, and Ridge regressor ^3^. The parameters were set to default values as found in the Sklearn library. Second, a simple feedforward neural network (NN) baseline is designed to directly map input features to throughput predictions without any explicit representation learning phase. Throughout this manuscript, the NN baseline consists of two components: a 5-layer linear encoder and a 5-layer linear decoder. These components are integrated into a single model, with ReLU activation functions applied between consecutive linear layers. The model is trained using mean squared error (MSE) loss. The third baseline is a Variational Autoencoder (VAE), which is composed of encoder-decoder architecture and trained to minimize the MSE Loss between the predicted and actual gene essentiality. The architecture, number of layers, training objective, and hyperparameters were adopted from this paper [34]. The model was trained to minimize the MSE loss between expression and essentiality. The code for the baselines can be found along with the paper materials.

## Footnotes

1. https://scikit-learn.org/1.5/modules/generated/sklearn.linear_model.LinearRegression.html

2. https://scikit-learn.org/1.5/modules/generated/sklearn.linear_model.Lasso.html

3. https://scikit-learn.org/1.5/modules/generated/sklearn.linear_model.Ridge.html

